# Enhancing antimicrobial resistance monitoring: Core Plasmid Multi-Locus sequence typing (cpMLST) with Oxford Nanopore sequencing technology (ONT)

**DOI:** 10.1101/2024.10.25.620091

**Authors:** Basil Britto Xavier, Anurag Kumar Bari, Tim T. Severs, Maria João Cardoso, Mykhailo Savin, Bhanu Sinha, John W. A. Rossen

## Abstract

**Background:** The spread of antimicrobial resistance (AMR) poses a significant threat to global public health, primarily driven by the horizontal gene transfer of resistance genes via plasmids. Understanding the dynamics of plasmid transmission within (intra) and between (inter) bacterial species is crucial for developing effective countermeasures. In this study, we utilize long-read sequencing technology to develop an innovative approach called core plasmid Multi-Locus Sequence Typing (cpMLST) and track the global plasmid transmission pathways.

**Methods:** We collected wastewater samples from six different hospitals in Germany. Within 24 hours of collection, we isolated extensively drug-resistant (XDR) bacteria using CHROMagar plates (CHROMagar, Paris, France). Species identification was done using MALDI-TOF MS (Bruker Daltonics, Bremen, Germany). Genomic DNA from the isolates was extracted using the PureLink Genomic DNA Mini Kit (Invitrogen, Carlsbad, USA) and subsequently sequenced using Oxford Nanopore Technologies (ONT) sequencing (Oxford Nanopore Technologies). We employed the SQK-RBK114.24 kit on an R10.4.1 flow cell, achieving a minimum of 250 Mb of raw data for each isolate with an accuracy of 98.9%. The reads were assembled using Unicycler, and the quality of the assemblies was assessed using QUAST and Bandage. The assembled genomes were then uploaded to the Center for Genomic Epidemiology (CGE) https://www.genomicepidemiology.org/ platform to determine incompatibility (Inc) types and antimicrobial resistance (AMR) genes. Additionally, we performed core plasmid analysis using chewBBACA v.3.3.10 to assess plasmid relatedness.

**Results:** The analysis confirmed the identification of various bacterial isolates. Among the most prevalent XDR isolate were *Enterobacter roggenkampii* (n=38), *Serratia marcescens* (n=12), *Klebseilla oxytoca* (n=11), *Klebseilla michiganensis* (n=8), and *Citrobacter freundii* (n=7). We conducted a cpMLST analysis to examine the highly prevalent plasmids and reveal evolutionary relationships and transmission patterns. Each isolate harbored different types of complete plasmids. Among those were the IncFII (n=45), IncHI (n=26), IncX3 (n=21), and other incompatibility groups. By focusing on shared Inc types among the bacterial species, their transfer within a single bacterial species (intra-species transfer) and between different species (inter-species transfer) were identified. The allelic differences ranged from 3-55 (IncHI), 2-32 (IncFII) and 2-24 (IncX3). Moreover, these plasmids, typically identified as smaller plasmids (IncX3), were larger than their commonly reported sizes. The analysis of plasmid dynamics across diverse bacterial hosts provides valuable insight into the mechanisms driving the spread of antimicrobial resistance, offering potential strategies for better controlling the dissemination of antimicrobial resistance and its transmission in clinical environments.

**Conclusion:** In conclusion, long-read sequencing technologies emerge as an essential tool for investigating the transmission dynamics of antimicrobial resistance genes among plasmids. By implementing cp-MLST, we anticipate achieving an unparalleled understanding of the global dissemination and evolutionary patterns of resistance-carrying plasmids. This approach offers superior resolution to conventional plasmid typing methods, potentially opening new avenues for innovative strategies to combat the escalating AMR crisis.

## Introduction

The global antimicrobial resistance (AMR) crisis, significantly driven by the widespread transmission of resistance plasmids, poses an alarming threat to modern healthcare systems. This crisis compromises our ability to effectively treat common infections, complicates routine medical procedures, and escalates healthcare costs. The World Health Organization (WHO) has identified AMR as one of the top ten global public health threats, highlighting the urgent need for coordinated strategies to mitigate this challenge (WHO, 2020). Plasmids are integral to the propagation of AMR among bacterial populations, playing a central role in this escalating health crisis. These extrachromosomal DNA molecules, which replicate independently of the bacterial chromosome, act as efficient vehicles for the horizontal transfer of genetic material, including antibiotic resistance genes (ARGs), across various bacterial species. The impact of plasmids on the spread of AMR is profound, as they enable the rapid dissemination of ARGs through horizontal gene transfer (HGT) mechanisms such as conjugation, transformation, and transduction. Conjugation facilitates the direct transfer of plasmids between bacterial cells via specialized structures, allowing for the swift spread of ARGs across both related and unrelated bacterial species (Carattoli, 2009; Partridge et al., 2018). Plasmids exhibit considerable diversity in size, ranging from a few kilobases to several hundred kilobases, and can carry multiple ARGs, conferring resistance to a broad spectrum of antibiotics. This multi-drug resistance capacity significantly amplifies their role in the AMR crisis (Forde & Zowawi, 2019). Moreover, plasmids often encode additional traits, including virulence factors and toxins, further contributing to bacterial evolution and adaptation (Fernández-López & de la Cruz, 2014). A variety of molecular typing methods have been developed to classify and track plasmids. These include PCR-based replicon typing (PBRT), which targets replicon sequences specific to different plasmid incompatibility groups; degenerate primer MOB typing (DPMT), which focuses on relaxase genes involved in plasmid mobilization; and plasmid multilocus sequence typing (pMLST), which offers finer resolution by analyzing multiple genetic loci on plasmids (A. Carattoli and H. Hasman., 2020; Lanza et al., 2014). Incompatibility (Inc) typing, based on the inability of plasmids with similar replication systems to coexist within the same cell, is also widely used (Orlek et al., 2017). However, these methods have limitations in terms of resolution and their ability to fully capture plasmid diversity. To better understand plasmid dynamics and AMR spread, advanced molecular typing methods and sequencing technologies are essential. Recent innovations in next-generation sequencing, particularly long-read sequencing (LRS) technologies like Oxford Nanopore Technologies (ONT), have significantly enhanced plasmid analysis. LRS provides higher resolution, enabling complete plasmid assemblies, improved detection of small plasmids, and more accurate mobilization typing, which is often limited by short-read sequencing technologies (Wyres & Holt, 2018; Wick et al., 2017). We propose that a cpMLST scheme can greatly improve plasmid typing. CpMLST offers a superior assessment of relatedness compared to traditional pMLST and Inc typing methods, providing a more comprehensive analysis by focusing on core plasmid genes. Moreover, cpMLST provides a standardized approach for plasmid classification across various studies and laboratories, enhancing the reproducibility and comparability of research findings (Li et al., 2018).

## Methods

### Isolate characterization

We collected wastewater samples from six different hospitals located across Germany. The purpose of this study was to isolate and identify strains of bacteria that exhibit extensive drug resistance (XDR). To ensure the integrity and accuracy of our results, we processed these samples within 24 hours of their collection. For the isolation of XDR strains, we utilized CHROMagar plates (CHROMagar, Paris, France). Following the isolation process, we proceeded with species identification using Matrix-Assisted Laser Desorption/Ionization-Time of Flight Mass Spectrometry (MALDI-TOF MS; Bruker Daltonics, Bremen, Germany).

### DNA extraction and LRS

Genomic DNA from the isolates was extracted using the PureLink Genomic DNA Mini Kit (Invitrogen, Carlsbad, USA) and used to prepare Oxford Nanopore Technologies (ONT) sequencing (Oxford Nanopore Technologies) libraries using Sequencing Kit (SQK-RBK114.24 kit). Sequencing was performed on the GridION X5 (v21.11.5) platform with R10.4.1 flow cells. Runs were conducted for 48-72 hours, and real-time base-calling was enabled resulting in a minimum of 250 Mb of raw data for each strain with an accuracy of 98.9% using Guppy (v6.4.2). Genome assembly was conducted using unicycler and Flye (v2.9) in parallel, which assembles long-read data into complete plasmid sequences. Post-assembly, Medaka (v1.7.1) was used for polishing to improve the accuracy of the consensus sequences derived from the raw reads.

### Core-plasmid MLST analysis

The assembled plasmids were annotated using Prokka (v1.14.6) (Seemann T. 2014), a comprehensive tool for identifying genes, coding sequences and other genetic elements on plasmids, including antibiotic resistance genes and virulence factors. Pangenome analysis was performed using Roary (v3.13.0) (Page JA., et al., 2015), focusing on the identification of core genes present in ≥95% of plasmids.

Core-plasmid Multi-Locus Sequence Typing (cpMLST) scheme was developed using chewBBACA (v3.3.10) (Silva M., et al., 2018). Briefly, identifying plasmids-specific core genes that are consistently present across plasmid genomes, followed by allele calling to determine which variant of each core gene is present in the query plasmid genomes. Then conducts a thorough comparison of these alleles across all plasmid sequences, calculating genetic distances based on the number of allelic differences observed between plasmids. Using these calculated distances, the clustering analysis to group genetically similar plasmids together, then minimum spanning trees were visualized using GrapeTree (v2.1) (Z Zhou et al., 2018) to illustrate the evolutionary relationships between plasmids.

## Results

### Genetic characterization

Sequencing results confirmed the identification of various bacterial isolates. The most prevalent XDR isolates were *Enterobacter roggenkampii* (n=38), *Serratia marcescens* (n=12), *Klebsiella oxytoca* (n=11), *Klebsiella michiganensis* (n=8) and *Citrobacter freundii* (n=7). We conducted a cpMLST analysis to examine the highly prevalent plasmids and reveal the evolutionary relationships and transmission patterns. Each isolate harbored different types of complete plasmids. Among those were the IncFII (n=45), IncHI (n=26), IncX3 (n=21), and other incompatibility groups. By focusing on shared Inc types among the bacterial species, their transfer within a single bacterial species (intra-species transfer) and between different species (inter-species transfer) were identified. The allelic differences ranged from 3-55 (IncHI2. Figure 1), 2-27 (IncN; Figure 2), 1-3 (IncX3; Figure 3) and 2-13 (IncFII Figure 4). Moreover, these plasmids, typically identified as smaller plasmids (IncX3) and were larger than their commonly reported sizes.

### IncHI2 plasmids

The IncHI2 and IncHI2A incompatibility groups were identified in the same plasmids found in *Klebsiella oxytoca, Enterobacter roggenkampii, Serratia marcescens*, and *Enterobacter asburiae*. These plasmids are primarily known for spreading antimicrobial resistance (AMR) genes, belong to antibiotic classes aminoglycosides, β-lactams, tetracyclines, sulfonamides, trimethoprim, including those that confer resistance to last-resort antibiotics carbapenems and colistin (Table 1). Plasmids can carry multiple replicons, which are sequences necessary for their replication and maintenance in bacterial cells. The presence of both IncHI2 and IncHI2A within the same plasmid suggests a multireplicon structure. This configuration can enhance plasmid stability and their ability to replicate across different bacterial hosts, thereby facilitating the spread of antimicrobial resistance genes. Additionally, this structure can undermine traditional Inc typing and plasmid multilocus sequence typing (pMLST). To address this challenge, core plasmid multilocus sequence typing (cpMLST) would be superior. The allelic differences in cpMLST range from 2 to 60 among different organisms and geographic origins (Figure 1).

**Figure 1.**
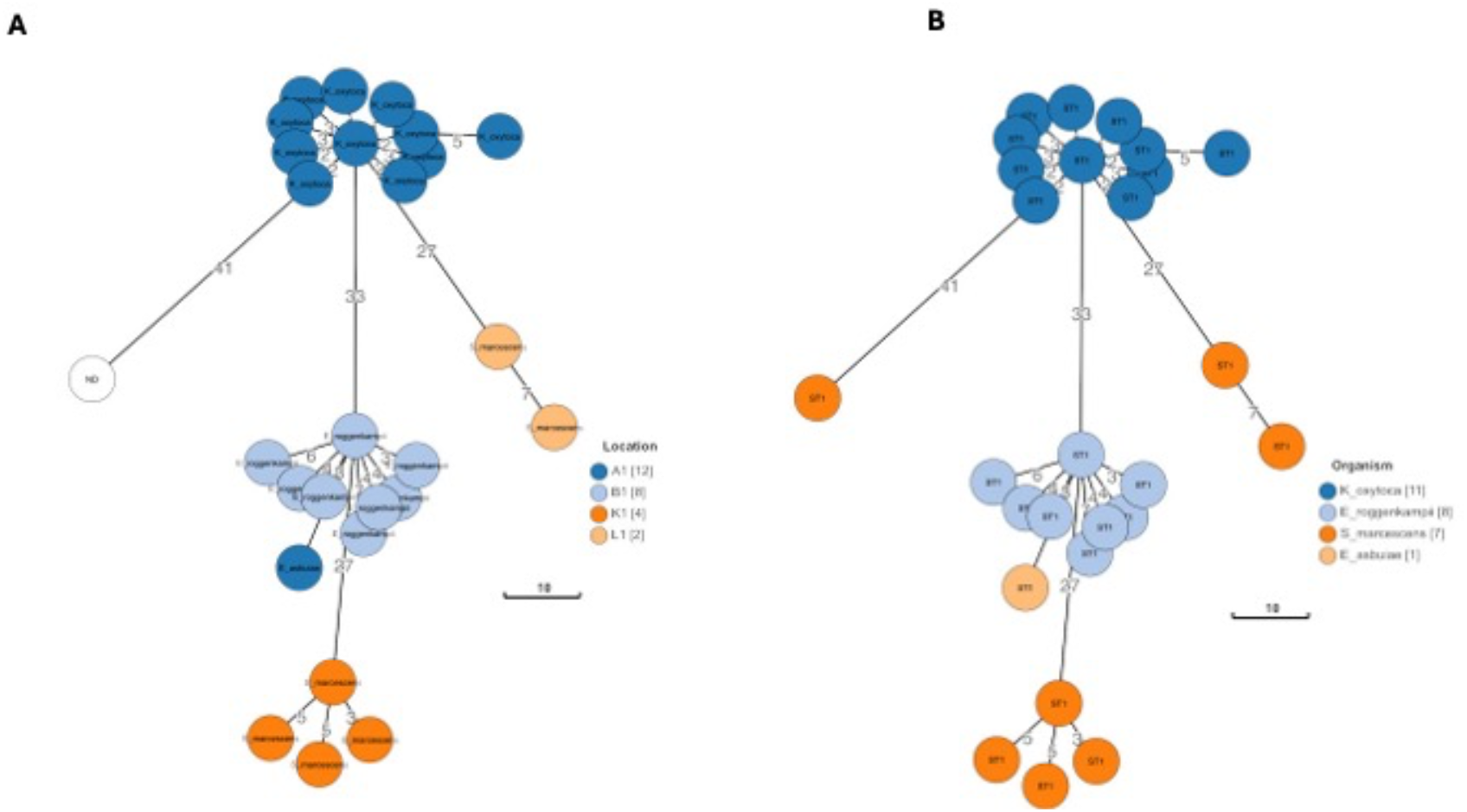
Minimum spanning trees of IncHI2 complete plasmids. The minimum spanning trees illustrate that complete plasmid sequences originating from different organisms yet belonging to the same pMLST type (ST1), exhibit diversity based on their geographic/locations (A) and organismal origins (B). Despite these differences, plasmids within the same population are closely linked, as indicated by minor allelic variations. The branches represent allelic differences at the core plasmid level. Organisms are as follows: *Klebsiella oxytoca* (blue), *Enterobacter roggenkampii* (light blue), *Serratia marcescens* (orange), and *Enterobacter asburiae* (yellow).

**Table 1:**
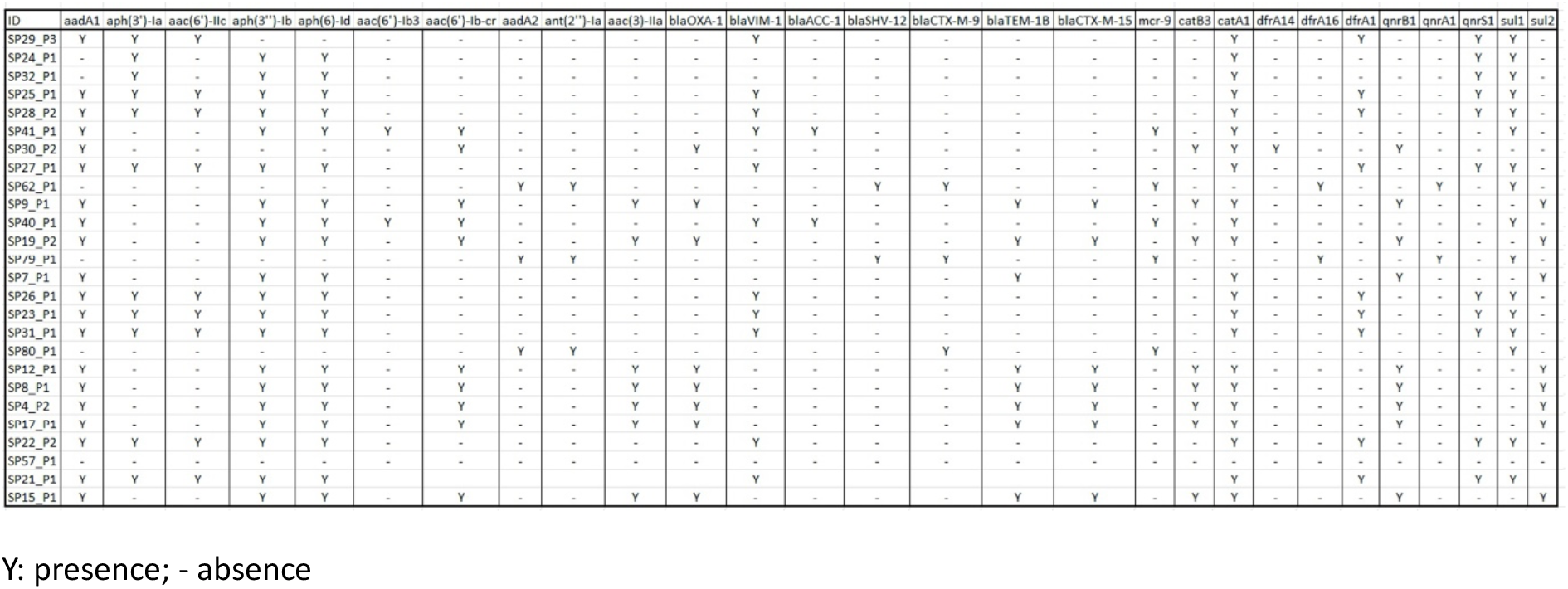
Resistome of IncHI2 plasmids.

### IncN Plasmids

The IncN incompatibility group has been identified in Enterobacter roggenkampii and Serratia marcescens. IncN plasmids are harbors antimicrobial resistance genes (ARGs), belong to antibiotic classes aminoglycosides, β-lactams, sulfonamides, trimethoprim, including those that confer resistance to last-resort antibiotic carbapenems (Table 2). It has been shown that cpMLST has enough discriminatory power to classify plasmids into the same or different sequence types (STs) on the same Inc type. This classification is based on comparing 101 core plasmid alleles rather than specific gene allele variants like the traJ, repN and korA loci. The allelic differences in the cpMLST among these analyzed plasmids range from 2 to 27 (Figure 2).

**Figure 2:**
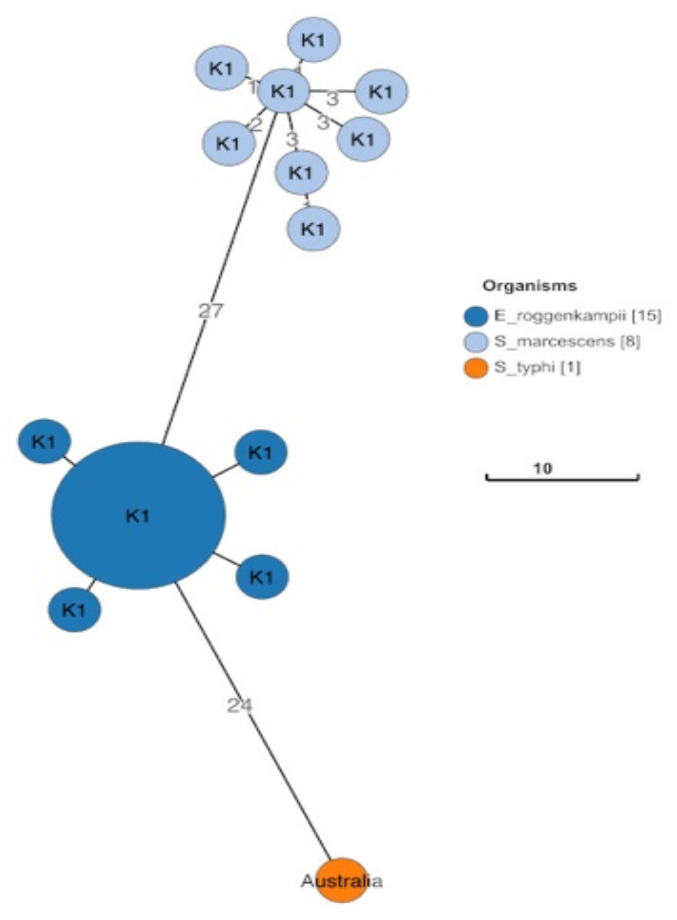
Minimum spanning tree of IncN complete plasmids. The minimum spanning tree illustrates that complete plasmid sequences originating from different organisms yet belonging to the same pMLST type (ST5), exhibit diversity based on their organismal origins. The size of the circle propitiates to the number of isolates. The branches represent allelic differences at the core plasmid level. Organisms are as follows: *Enterobacter roggenkampii* (blue), *Serratia marcescens* (light blue).

**Table 2:** Resistome of IncN plasmids.

| ID | aac(6')Ib3 | aadA1 | aac(6')Ibcr | aac(3)IId | aph(3'')Ib | aph(6)Id | tet(A) | qnrB2 | sul1 | blaKPC2 | blaVIM1 | blaOXA1 | blaTEM1B | mph(A) | catB3 | ARR3 | dfrA19 |
| --- | --- | --- | --- | --- | --- | --- | --- | --- | --- | --- | --- | --- | --- | --- | --- | --- | --- |
| SP8Y_P2 | Y | Y | Y | - | - | - | Y | Y | * | - | Y | - | - | - | - | - | - |
| SP78_P2 | Y | Y | Y | - | - | - | Y | Y | * | - | Y | - | - | - | - | - | - |
| SP76_P2 | Y | Y | Y | - | - | - | Y | Y | * | - | Y | - | - | - | - | - | - |
| SP65_P2 | Y | Y | Y | - | - | - | Y | Y | * | - | Y | - | - | - | - | - | - |
| SP52_P2 | Y | Y | Y | - | - | - | Y | Y | * | - | Y | - | - | - | - | - | - |
| SP63_P3 | - | - | - | Y | Y | Y | - | - | - | Y | - | - | Y | - | - | - | - |
| SP73_P3 | Y | Y | Y | - | - | - | Y | Y | * | - | Y | - | - | - | - | - | - |
| SP68_P3 | Y | Y | Y | - | - | - | Y | - | Y | - | Y | - | - | - | - | - | - |
| SP80_P3 | - | - | - | Y | Y | Y | - | - | - | Y | - | - | Y | - | - | - | - |
| SP55_P2 | Y | Y | Y | - | - | - | Y | Y | * | - | Y | - | - | - | - | - | - |
| SP57_P3 | - | - | - | Y | Y | Y | - | - | - | Y | - | - | Y | - | - | - | - |
| SP79_P3 | - | - | - | - | Y | Y | - | - | - | - | - | - | - | - | - | - | - |
| SP7Y_P2 | Y | Y | Y | - | - | - | Y | Y | * | - | * | - | - | - | - | - | - |
| SP77_P2 | Y | Y | Y | - | - | - | Y | Y | * | - | Y | - | - | - | - | - | - |
| SP58_P2 | - | - | Y | Y | Y | Y | - | Y | * | Y | - | Y | Y | Y | Y | Y | Y |
| SP60_P2 | - | - | Y | Y | Y | Y | - | Y | * | Y | - | Y | Y | Y | Y | Y | Y |
| SP74_P3 | Y | Y | Y | - | - | - | Y | Y | * | - | Y | - | - | - | - | - | - |
| SP59_P2 | Y | Y | Y | - | - | - | Y | - | Y | - | Y | - | - | - | - | - | - |
| SP70_P2 | Y | Y | Y | - | - | - | Y | Y | * | - | Y | - | - | - | - | - | - |
| SP64_P2 | Y | Y | Y | - | - | - | Y | Y | * | - | Y | - | - | - | - | - | - |
| SP62_P3 | - | - | - | Y | Y | Y | - | - | - | Y | - | - | Y | - | - | - | - |
| SP75_P3 | Y | Y | Y | - | - | - | Y | Y | * | - | Y | - | - | - | - | - | - |
| SP79_P5 | - | - | - | Y | Y | Y | - | - | - | Y | - | - | Y | - | - | - | - |

Y: presence; - absence

### IncX Plasmids

IncX plasmids, primarily found in *Enterobacterales*, are categorized into subgroups like IncX1 and IncX2 based on Inc typing. Their diversity is highlighted by their ability to carry various resistance genes, including those against carbapenems and β-lactams (Table 3). However, the absence of a standardized plasmid multilocus sequence typing (pMLST) scheme complicates their classification and tracking, hindering epidemiological and evolutionary studies. We have used core plasmid multilocus sequence typing (cpMLST), which involves comparing numerous loci across plasmid genomes to identify core genes common among them. For IncX plasmids, 212 loci were compared, with 36 identified as core loci (Figure 3), providing a high-resolution framework for understanding their genetic diversity and evolutionary relationships. This method allows for a detailed understanding of the genetic structure and diversity of IncX plasmids by focusing on conserved core genes, offering insights into the evolutionary pressures and mechanisms driving the spread of antibiotic resistance.

**Figure 3.**
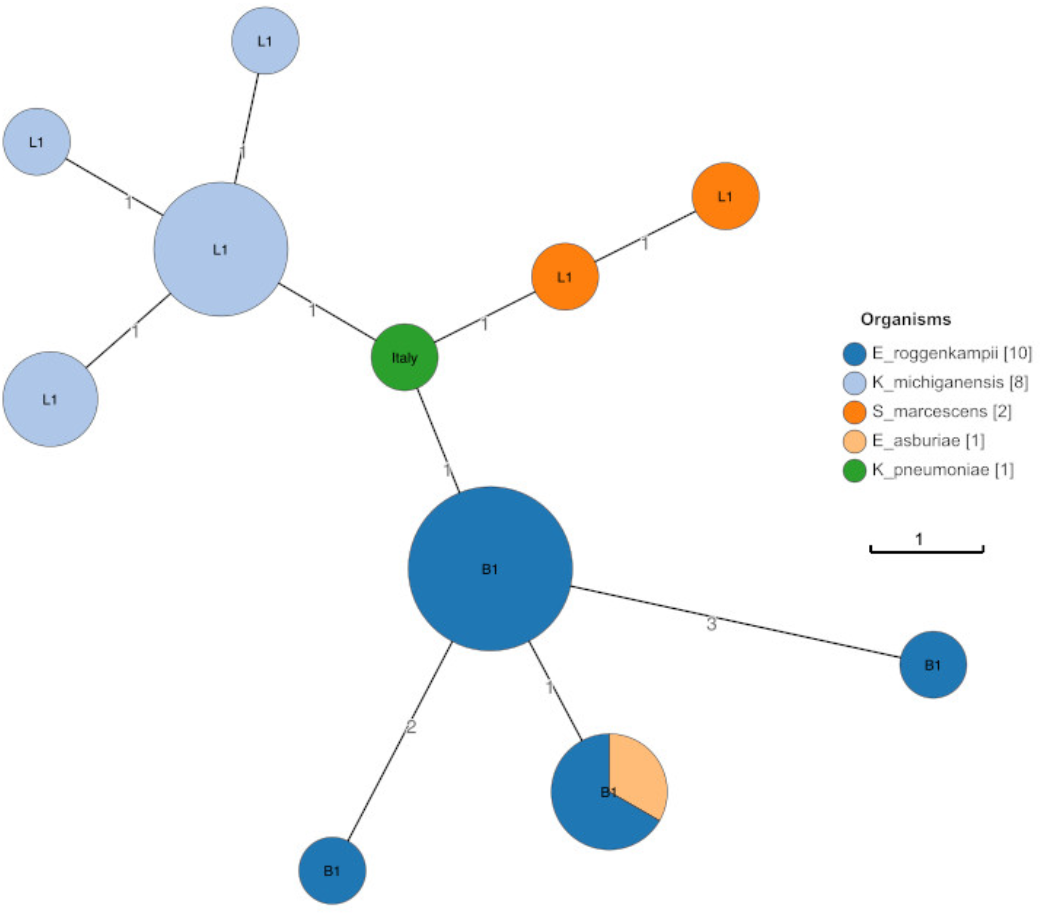
Minimum spanning tree of IncX complete plasmids. The minimum spanning tree illustrates that complete plasmid sequences originating from different organisms yet belonging to the same pMLST type (ST1), exhibit diversity based on their geographic/locations and organismal origins. Despite these differences, plasmids within the same population are closely linked, as indicated by minor allelic variations. The branches represent allelic differences at the core plasmid level. Organisms are as follows: *Enterobacter roggenkampii* (blue), *Klebsiella michiganensis* (light blue), *Serratia marcescens* (orange), and *Enterobacter asburiae* (yellow) [*Klebsiella pneumoniae* (green].

**Table 3:** Resistome of IncX plasmids.

| ID | blaNDM-10 | blaNDM-18 | blaNDM-1 | blaNDM-24 | blaKPC-2 | blaSHV-182 | blaSHV-12 |
| --- | --- | --- | --- | --- | --- | --- | --- |
| SP7_P3 | Y | - | - | - | - | - | - |
| SP47_P4 | - | - | - | - | - | - | Y |
| SP15_P3 | - | Y | - | - | - | - | - |
| SP43_P6 | - | - | - | - | - | - | - |
| SP17_P3 | - | Y | - | - | - | - | - |
| SP8_P3 | - | - | Y | - | - | - | - |
| SP42_P2 | - | - | - | - | - | - | Y |
| SP13_P2 | - | - | - | - | - | - | - |
| SP19_P5 | - | - | - | - | - | - | - |
| SP30_P3 | - | - | Y | - | - | - | - |
| SP12_P2 | - | - | Y | - | - | - | - |
| SP45_P4 | - | - | - | - | - | - | Y |
| SP44_P4 | - | - | - | - | - | - | Y |
| SP51_P4 | - | - | - | - | - | - | Y |
| SP9_P3 | - | - | Y | - | - | - | - |
| SP16_P2 | - | - | - | Y | - | - | - |
| SP4_P4 | - | - | - | - | - | - | - |
| SP46_P5 | - | - | - | - | - | - | Y |
| SP48_P3 | - | - | - | - | Y | - | Y |
| SP49_P4 | - | - | - | - | Y | - | Y |
| SP50_P4 | - | - | - | - | - | - | Y |

Y: presence; - absence

### IncFII plasmids

The IncFII incompatibility groups were identified in *Enterobacter roggenkampii, Klebsiella oxytoca, Citrobacter farmeri, Serratia marcescens, Klebsiella michiganensis*, and *Enterobacter soli*. IncF plasmids are divided into subgroups such as IncFII and IncFIB, among other variants, based on Inc typing. Their diversity is underscored by their capacity to carry various resistance genes, including those antibiotic classes aminoglycosides, tetracyclines, carbapenems, sulfonamides, sulfonamides and β-lactams (Table 4). However, the lack of a standardized and harmonized Inc typing and plasmid multilocus sequence typing (pMLST) scheme complicates their classification and tracking, posing challenges for epidemiological and evolutionary studies. To address this, we performed core plasmid multilocus sequence typing (cpMLST) to determine the relatedness of plasmids belonging to the broader IncF compatibility group from different geographical and organismal origins. We successfully determined the relatedness among these plasmids, finding core plasmid allelic variations ranging from a minimum of 1 to a maximum of 13 when compared with 308 allelic variations across the entire plasmid (Figure 4).

**Figure 4.**
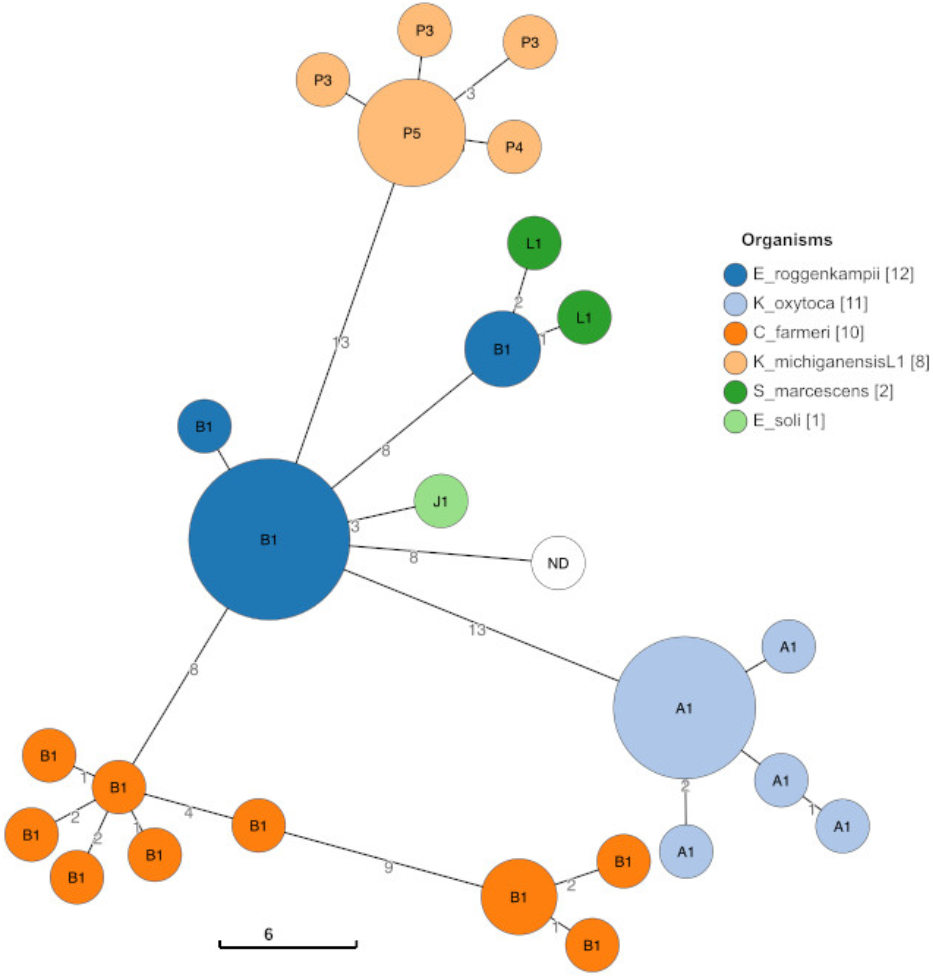
Minimum spanning tree of IncFII complete plasmids. The minimum spanning tree illustrates that complete plasmid sequences originating from different organisms yet belonging to the different pMLST type and Inc type, exhibit diversity based on their geographic/locations and organismal origins. Despite these differences, plasmids within the same population are closely linked, as indicated by minor allelic variations. The branches represent allelic differences at the core plasmid allelic differences. Organisms are as follows: Enterobacter *rogenkompii* (dark blue), *Klebsiella oxytoca* (light blue), *Citrobacter farmeri* (orange), *Klebsiella michiganensis* (light orange), *Serratia marcescens* (green), and *Enterobacter soli* (light green).

**Table 4:** Resistome of IncF plasmids from this study.

| ID | aac(6')-Ib3 | blaOXA-2 | aac(6')-Ib-cr | aadA1 | aac(6')-Ib | rmtC | blaTEM-1A | blaOXA-9 | blaSHV-12 | blaNDM-1 | blaVIM-1 | sul1 | qnrS1 | tet(A) |
| --- | --- | --- | --- | --- | --- | --- | --- | --- | --- | --- | --- | --- | --- | --- |
| SP19_P4 | - | - | - | - | - | - | - | - | - | - | - | - | - | - |
| SP28_P4 | - | - | - | - | - | - | - | - | - | - | - | - | - | - |
| SP48_P1 | Y | Y | Y | - | - | - | - | - | - | - | - | - | - | - |
| SP10_P3 | - | - | Y | Y | Y | Y | Y | Y | Y | Y | - | Y | - | - |
| SP23_P2 | - | - | - | - | - | - | - | - | - | - | - | - | - | - |
| SP4_P3 | - | - | - | - | - | - | - | - | - | - | - | - | - | - |
| SP15_P2 | - | - | - | - | - | - | - | - | - | - | - | - | - | - |
| SP5_P5 | - | - | - | - | - | Y | - | - | - | Y | - | Y | - | - |
| SP14_P2 | - | - | - | - | - | Y | - | - | - | Y | - | Y | - | - |
| SP13_P1 | - | - | - | - | - | - | - | - | - | - | - | - | - | - |
| SP20_P4 | - | - | Y | Y | Y | - | Y | Y | Y | - | - | - | - | - |
| SP11_P3 | - | - | - | - | - | Y | - | - | - | Y | - | Y | - | - |
| SP25_P2 | - | - | - | - | - | - | - | - | - | - | - | - | - | - |
| SP24_P2 | - | - | - | - | - | - | - | - | - | - | - | - | - | - |
| SP20_P3 | - | - | - | - | - | Y | - | - | - | Y | - | Y | - | - |
| SP2_P3 | - | - | - | - | - | - | - | - | - | Y | - | - | - | - |
| SP8_P2 | - | - | - | - | - | - | - | - | - | - | - | - | - | - |
| SP17_P2 | - | - | - | - | - | - | - | - | - | - | - | - | - | - |
| SP16_P1 | - | - | - | - | - | - | - | - | - | - | - | - | - | - |
| SP45_P3 | - | - | - | - | - | - | - | - | - | - | - | - | Y | - |
| SP42_P3 | - | - | - | - | - | - | - | - | - | - | - | - | Y | - |
| SP31_P2 | - | - | - | - | - | - | - | - | - | - | - | - | - | - |
| SP11_P4 | - | - | Y | Y | Y | - | Y | Y | Y | - | - | - | - | - |
| SP3_P4 | - | - | - | - | - | - | - | - | - | - | - | - | - | - |
| SP50_P5 | - | - | - | - | - | - | - | - | - | - | - | - | Y | - |
| SP29_P1 | - | - | - | - | - | - | - | - | - | - | - | - | - | - |
| SP7_P2 | - | - | - | - | - | - | - | - | - | - | - | - | - | - |
| SP12_P3 | - | - | - | - | - | - | - | - | - | - | - | - | - | - |
| SP44_P3 | - | - | - | - | - | - | - | - | - | - | - | - | Y | - |
| SP49_P2 | Y | Y | Y | - | - | - | - | - | - | - | - | - | - | - |
| SP22_P1 | - | - | - | - | - | - | - | - | - | - | - | - | - | - |
| SP46_P4 | - | - | - | - | - | - | - | - | - | - | - | - | Y | - |
| SP26_P2 | - | - | - | - | - | - | - | - | - | - | - | - | - | - |
| SP21_P2 | - | - | - | - | - | - | - | - | - | - | - | - | - | - |
| SP32_P2 | - | - | - | - | - | - | - | - | - | - | - | - | - | - |
| SP1_P2 | - | - | Y | Y | Y | Y | Y | Y | Y | Y | - | Y | - | - |
| SP51_P3 | - | - | - | - | - | - | - | - | - | - | - | - | Y | - |
| SP43_P5 | - | - | - | - | - | - | - | - | - | - | - | - | Y | - |
| SP5_P6 | - | - | Y | Y | Y | - | Y | Y | Y | - | - | - | - | - |
| SP27_P2 | - | - | - | - | - | - | - | - | - | - | - | - | - | - |
| SP33_P3 | Y | - | Y | Y | - | - | - | - | - | - | Y | Y | - | Y |
| SP47_P3 | - | - | - | - | - | - | - | - | - | - | - | - | Y | - |
| SP18_P2 | - | - | - | - | - | - | - | - | - | - | - | - | - | - |
| SP9_P2 | - | - | - | - | - | - | - | - | - | - | - | - | - | - |

Y: presence; - absence

## Discussion

In this cpMLST analysis, using multiple core genes significantly enhances phylogenomic resolution compared to traditional methods like Inc typing and pMLST. This approach is like bacterial core genome level comparisons, such as core genome MLST (cgMLST), which utilize tools like chewBBACA (Silva M., et al., 2018) and Ridom SeqSphere+. The Validated Bacterial Core Genes (VBCG) set, consisting of 20 genes, offers improved discrimination between species and strains while maintaining alignment with established phylogenies. This method effectively addresses the variability issues associated with individual genes, providing a more robust evolutionary signal. Recent advancements in LRS technologies, such as those from ONT, have further enhanced plasmid analysis by enabling the full reconstruction of plasmid sequences. This capability is crucial for understanding plasmid diversity and evolution, particularly in relation to the spread of ARGs. ONT sequencing can detect repeats and other structural variations, including insertions, deletions, and single nucleotide variants, which are essential for tracking the spread of AMR. Here we developed cpMLST schemes to improve plasmid typing accuracy by identifying core genes present in a high percentage of plasmids and selecting stable and variable genes to define STs. The cpMLST approach offers superior discriminatory power compared to pMLST and Inc typing and aligns well with whole-plasmid phylogenies. Overall, integrating ONT LRS with cpMLST provides a comprehensive approach to plasmid typing, offering detailed insights into bacterial genetics and the spread of AMR. As technology advances and costs decrease, this method is poised to become a vital tool in microbial genomics and epidemiology.

While LRS combined with cpMLST offers significant advantages, this study faces several limitations that must be addressed. First, LRS is associated with higher error rates compared to short-read sequencing, which could influence the determination of allelic differences at the core plasmid level. Although advances in ONT sequencing chemistries and analysis methods are improving accuracy, this remains a concern. Additionally, bioinformatics challenges persist, as base calling and analyzing long-read data require specialized tools and significant computational resources. Moreover, LRS often requires larger quantities and higher quality DNA input, which can be difficult to obtain from certain sample types. Furthermore, publicly available cpMLST schemes for plasmid typing are still under development and may not yet be as comprehensive as those available for other typing methods such as pMLST and Inc typing. Existing databases, built primarily with short-read data, may also limit their applicability to LRS data. Despite these limitations, the benefits of LRS combined with cpMLST for studying bacterial plasmids remain substantial, and ongoing technological advancements are expected to address these challenges.

## In conclusion

specific cpMLST schemes for every plasmid type can be developed to trace and track plasmids within populations with high resolution and broad coverage of plasmid diversity. This is essential for understanding the spread of ARGs, contributing to a deeper understanding of AMR transmission dynamics and supporting more effective strategies to combat (the spread of) AMR.

## Acknowledgment

This study was partly supported by DRAIGON: Diagnosing infections with Multi-Drug-Resistant Microorganisms using AI-Powered Genomic Antibiotic Susceptibility Prediction from Long-read sequencing data project funded by the European Union. Views and opinions expressed are, however, those of the author(s) only and do not necessarily reflect those of the European Union or European Health and Digital Executive Agency (HADEA). Neither the European Union nor the granting authority can be held responsible for them. This project has received funding from the European Union’s Horizon Europe research and innovation programme under grant agreement No 101137383. Partly the study also supported by SPOWAR: Sustainable Protection of WAter Resources funded by the European Union and the Interreg partners as part of the Germany-Netherlands Interreg program: INTERREG VI A, No 24035.

